# Modelling cognitive outcomes in the UK Biobank: education, noradrenaline and frontoparietal networks

**DOI:** 10.1101/2023.08.31.555645

**Authors:** Laura Bravo-Merodio, Jackie A. Williams, Dominic Russ, Georgios Gkoutos, Meadhbh B. Brosnan, Mark A. Bellgrove, Magdalena Chechlacz

## Abstract

Education is often used as a surrogate measure of so called cognitive reserve (CR) benefiting cognitive functioning in later years. In line with Robertson’s theory we tested here a hypothesis that education acting on the noradrenergic system strengthen the right fronto-parietal networks to facilitate CR and maintain cognition throughout the lifetime. We used machine learning and mediation analysis to model interactions between neurobiological features (genetic variants in noradrenergic signalling, structural and functional fronto-parietal connectivity) and education (proxy of CR) on cognitive outcomes (general cognitive ability score) in the UK Biobank cohort. We show that: (1) interactions between education and neurobiological variables better explain cognitive outcomes than either factor alone; (2) among the neurobiological features selected using variable importance testing, measures of right fronto-parietal connectivity are the strongest mediators of the effect of education on cognitive outcomes. Our findings offer novel insights into neurobiological basis of CR by pointing to between-networks connectivity, representing connections linking the default mode network with the right fronto-parietal network as the key facilitator of CR.

## INTRODUCTION

The global demographic shift over the last few decades leading to increased proportion of older adults resulted in steadfastly raising numbers of people at risk of cognitive decline and dementia. This has long-lasting impact not only on the individual quality of life but also major economic impact on the societies worldwide (Livingston et al., 2020). While the ageing process is inevitable and comes with expectation of worsening in cognitive functioning, the literature examining cognitive ageing points to a striking heterogeneity in the rate of age-related deterioration in cognitive decline. It means that some older adults retain high levels of mental capacity, some undergo a gradual drop in cognitive functioning, and finally some experience a sharp cognitive decline (dementia) hindering their ability to undertake basic daily activities and affecting quality of life (Rapp and Amaral 1992, Hayden et al., 2011, Norton et al., 2014; Cabeza et al., 2018; Stern et al., 2020). To keep up with the ageing population and to enable advances in medical/social care a better understanding of factors affecting these different cognitive outcomes is urgently needed (Prince et al., 2015; Livingstone et al., 2020).

Cognitive ageing is a highly complex process caused by a convergence of multiple neurobiological and neurophysiological changes influenced by environmental and genetic factors as well as gene-environment interactions (e.g., Harris and Deary, 2011; Davies et al., 2015; Lu et al., 2016; Tucker-Drob et al., 2014, Papenberg et al., 2016). Among environmental influences, sociodemographic and lifestyle factors seem to play a critical role (Stern et al., 2020; Cabeza et al., 2018). It is well evidenced that the cognitive ageing is a result of structural and functional brain changes causing deterioration of cognitive performance. However, the interpretation of multifactorial influences on age-related changes in neural networks underlying the differential trajectories of cognitive ageing presents an ongoing challenge.

There is a growing evidence that older adults, who have engaged across lifespan in cognitively and/or socially enriched environments exhibit greater resilience to cognitive decline and maintain better brain health and cognitive function in later year. The concept of cognitive reserve (CR) has been proposed to account for at least some the observed heterogeneity in healthy (non-dementia) cognitive ageing as well as susceptibility to dementia, pathological decline (Stern et al., 2020; Cabeza et al., 2018). Within this framework, CR is measured by proxies of life experiences, in particular education but also occupation and leisure activities (Cabeza et al., 2018; Valenzuela and Sachdev, 2006a,b; Opdebeeck et al., 2015; Nucci et al., 2012).

While the concept of CR offers appealing framework for understanding cumulative neural changes due to lifelong environmental influences, including specifically socio-behavioural factors, and their interplay with genetic variability, yet to date few theoretical and experimental attempts have been made in understanding the neurobiological origins of cognitive reserve in healthy ageing (Cabeza et al., 2018; Robertson et al., 2014, Brosnan et al., 2018, Brosnan et al., 2022; Shalev at al., 2020; Plini et al., 2021). Here, we used one such theoretical framework (Robertson’s theory; Robertson et al., 2014) of CR to test the potential of machine learning methods to tease apart combinatorial effects of education (as CR proxy) and genetic influences on brain networks and cognitive outcomes (heterogeneity in cognitive function) in a large cross-sectional ageing cohort, the UK Biobank study. As UK Biobank has been created as a large-scale epidemiological resource with extensive sociodemographic questionnaires, physical measures, medical records, neuroimaging, and genetic data from middle and old age participants (Sudlow et al., 2015), it enables to tease apart multifactorial influences on cognitive ageing, using data science approaches on a scale not feasible before.

Robertson’s theory of CR proposes that life experiences acting on the noradrenergic system strengthen the right fronto-parietal networks to facilitate cognitive reserve and maintain cognition throughout lifetime (Robertson et al., 2013; Robertson et al., 2014). This theory combines Stern’s observations on the effects of education in Alzheimer’s patients with animal research. Animal studies provide evidence that across the life-span neural networks are heavily subjected to regulatory influences of neuromodulators (dopamine, noradrenaline, serotonin, and acetylcholine). These neuromodulators are thought to maintain functional dynamics among large-scale brain networks and to optimize cognitive performance by signalling environmental inputs (for review see Avery and Krichmar, 2017). In line with animal research Robertson’s theory stipulates that life experiences trigger continues engagement of several core cognitive process, all of which functionally rely on locus coeruleus— noradrenergic system and right fronto-parietal networks (Robertson et al., 2013; Robertson et al., 2014). A recent study linking cognitive performance and brain health in both non-demented older adults and Alzheimer’s patients to the volumetric estimates of locus coeruleus has provided compelling evidence that indeed the noradrenergic system might underpin the CR (Plini et al., 2021). There is also evidence in support of right lateralised cognitive reserve network (van Loenhoud et al., 2017; Shalev et., 2020, Brosnan et la., 2018; Brosnan et al., 2022). For example, our group has previously shown that life experiences mitigate age-related cognitive deficits by (i) preserving grey matter withing the right fronto-parietal regions (Shalev et al., 2020) and (ii) offsetting age-related axonal dispersion within the right fronto-parietal white matter (Brosnan et al., 2022). Interestingly, we previously demonstrated that both the composite cognitive reserved index measure (based on education, occupation and cognitively stimulating leisure activities) and the education alone have an offsetting effect on age-related changes in attention function by preserving grey matter within the right fronto-parietal networks (Shalev et al., 2020). It should be noted that none of the previous studies examining neurobiological basis of Robertson’s theory of CR simultaneously tapped into the noradrenergic system and the right fronto-parietal networks, which current study addresses.

Growing evidence suggests that genetic variations, leading to either elevated or decreased levels of neuromodulators have impact on the functional dynamics of the neural networks, and underlie inter-individual variability in cognitive abilities across human lifetime (Lindenberger et al., 2008; Papenberg et al., 2015). Thus, along these lines Robertson’s theory of CR presents an interesting framework for understanding mechanism of cumulative socioeconomic and genetic influences on functional neural networks and whether such interplay contributes to CR. It enables to stipulate that genetic variability, enhancing neuromodulation across lifespan, and strengthening the right fronto-parietal networks in response to life experiences (e.g., education) could be advantageous in offsetting age-related cognitive decline. To test such proposal, we used here machine learning approaches, previously successfully applied by our group in large epidemiological studies (Bravo-Merodio et al., 2019; COVIDSurg Collaborative et al., 2021) to model interactions between education and neurobiological features (measures of structural and functional frontoparietal connectivity and genetic variants in noradrenergic signalling) on cognitive outcomes in the ageing UK Biobank cohort.

## METHODS

### Participants

We used data from the UK Biobank, a prospective epidemiological cohort study with over 500,000 participants 40–69 years of age at recruitment (total of 502,505 participants), who underwent wide-ranging phenotypic and genotypic characterisation (Sudlow et al., 2015). For the purpose of the current study, we employed (1) genomic, (2) demographic data from the initial baseline visit at recruitment between 2006 and 2010 (i.e., initial assessment visit), and (3) cognitive and (4) multimodal neuroimaging data acquired between 2014 and 2020 (i.e., imaging visit; the UK Biobank Brain Imaging Cohort; Miller et al., 2016; Alfaro-Almagro et al., 2018). At the time of the study, multimodal neuroimaging data release included 48,561 participants, and so modelling was performed with these datasets only. All the analyses were conducted under the UK Biobank application numbers 29447 and 31224. All UK Biobank participants provided written informal consent in accordance with approved ethics protocols (REC reference number 11/NW/0382).

### Demographic information and cognitive data

Basic demographic information including month and year of birth, sex and education was recorded as part of information acquired at recruitment or as part of touchscreen questionnaire completed during the baseline assessment visit. The full details of the touchscreen questionnaire and all procedures are provided on the UK Biobank website (https://biobank.ndph.ox.ac.uk/ukb). The month and year of birth alongside the date of attending imaging visit (MRI scanning session) were used to calculate participants age. For the purpose of the current study, we used sex as recorded at recruitment (information derived from central registry i.e., as recorded by NHS or if not available, self-reported information was used). Education variable was determined based on self-reported age of completing full time education. This variable does not capture whether someone completed undergraduate or graduate degree. The variable captures instances where there was a break in education with intention to return but does not capture return to full time studies later in life. Most UK Biobank participants provided this information during baseline visit by answering a question “At what age did you complete your continuous full time education?”, with those reporting they “Never went to school” having a 0 assigned.

However, if this information was not captured during the baseline assessment (instances of a missing answer as opposed to participant answering “Prefer not to answer” (2760) or “Do not know” (3454), it was gathered during the subsequent visits. Notwithstanding, we identified a large portion of participants 46% (or 22,334 out of 48,705 participants) with this information still missing (Supplementary Table 1). After ascertaining that this information was Missing At Random (MAR) (Supplementary Table 1, Supplementary Figure 1) and given its use as proxy of cognitive reserve, participants with missing education information were deleted from our main analyses. Subsequently as explained below, the same analyses were performed using imputed data.

Cognitive assessment was completed on the same day as attending imaging visit and entailed completing several simple cognitive tests administered via touchscreen. The UK Biobank cognitive test battery consists of 13 different tests as detailed in the UK Biobank showcase (https://biobank.ndph.ox.ac.uk/showcase/label.cgi?id=100026). However, some of these tests were only used during piloting stage of the UK Biobank study and some tests added or modified for the subsequent recruitment phases and/or subsequent visits. Additional information about UK Biobank cognitive tests, alongside their validity and reliability assessment can be found in the previously published papers (Sudlow et al., 2015; Fawns-Ritchie and Deary, 2020). For the purpose of the current study we used a subset of measures of cognitive performance on 10 different tests measuring memory, reasoning, executive function and processing speed: (1) two symbol matching card game measuring reaction time calculated as the mean response time to correctly identify matching pairs of cards, (2) digit span test with a maximum recall set to twelve digits measuring a short term numeric memory calculated as the maximum number of correctly recalled digits, (3) fluid intelligence test with thirteen questions measuring reasoning and problem solving based on a total number of correct answers (i.e., fluid intelligence score), (4) version A of Trial Making test assessing visuospatial attention based on a time required to complete numeric path by correctly connecting 25 consecutive numbers; (5) version B of Trial Making test assessing visuospatial attention and task switching based on time required to complete alphanumeric path by connecting alternating numbers (1-13) and letters (A-L), (6) matrix patterns test assessing abstract reasoning based on number of correctly (out of fifteen) solved puzzles, (7) tower rearranging test measuring executive function based on the number of correctly solved puzzles (i.e., correctly guest number of moves required to re-arrange hoops to match a target “tower” image), (8) symbol digit substitution test measuring multiple processing abilities (attention, associative learning, visual processing) based on number of correctly identified symbol-digit matches, (9) paired associate learning test measuring number of correctly matched (out of twelve) word pairs, and (10) pairs-matching test measuring visual memory (number of errors = incorrect matches) tested based on assessing memorized positions of six pairs of cards.

### MRI data

The neuroimaging datasets available for the UK Biobank Brain Imaging Cohort (i.e., participants who completed imaging visit) include a set of imaging derived phenotypes (IDPs) i.e., various measures derived from different magnetic resonance imaging (MRI) modality-specific analyses (Alfaro-Almagro et al., 2018). The full information about the MRI data acquisition and data processing pipelines applied to the data provided by the UK Biobank is available online (https://biobank.ndph.ox.ac.uk/ukb/ukb/docs/brain_mri.pdf) and in previously published primary UK Biobank methods papers (Miller et al., 2016; Alfaro-Almagro et al., 2018). For the purpose of the current study we selected a set of IDPs representing structural and functional fronto-parietal connectivity measures derived from diffusion MRI (dMRI) and resting state functional MRI (rs-fMRI) respectively. Specifically, we used dMRI IDPs reflecting microstructural properties of white matter pathways, which were derived from neurite orientation and dispersion imaging (NODDI) model and representing intra-cellular volume fraction (ICVF) as a measure of neurite packing density, and the orientation dispersion index (ODI) as a measure of dispersion of neurites (an estimate of fiber coherence) for three fronto-parietal white matter pathways, superior longitudinal fasciculus (SLF), inferior-fronto-occipital fasciculus (IFOF) and the forceps minor of the corpus callosum. In addition, we used rs-fMRI IDPs reflecting brain functional connectivity, which were generated by (1) carrying out group independent component analysis (ICA) parcellations with dimensionality (the number of distinct ICA components identified based on temporal patterns of spontaneous fluctuations in brain activity) set at 25, (2) removal of noise components resulting in 21 components representing separate resting state networks (functional nodes), and (3) estimating 21×21 partial correlation matrices, which represent direct connections (edges) between pairs of ICA components (nodes), see FSLNets user guide (https://fsl.fmrib.ox.ac.uk/fsl/fslwiki/FSLNets). For the purpose of the current study, we have selected 6 networks of interests (left and right fronto-parietal network, executive control network and 3 subsystems of the default mode network: core (cDMN), dorsomedial prefrontal (dmDMN) and medial temporal (mtDMN); 6 networks representing frontoparietal connectivity and functionally related to core cognitive processes as per Robertson’s theory; identified based on previously published papers (Smith et al., 2009; Laird et al., 2009; Shirer et al., 2012, Dixon et al., 2017); see Supplementary Figure 2) and connection (edges) between them were entered into the analysis (total of 15 different edges). The 25 components group-ICA spatial maps and connection edges are available online (http://www.fmrib.ox.ac.uk/ukbiobank).

### Missing value imputation

Given education was used here as proxy of cognitive reserve, we first deleted all participants with missing information (22,334 participants) (Supplementary Table 1 and Supplementary Figure 1). From this dataset (26,227 participants and 39 variables spanning demographic data, cognitive tests and multimodal neuroimaging data), two different approaches were taken in order to deal with missing data information. First, we performed a complete case analyses, deleting all patients with missing value information, after having seen no visible pattern in previous analysis and determining missing values were missing at random (MAR) (Supplementary Figure 3). The final dataset consisted of 12,076 participants. The second approach consisted in imputing missing value data (20.2%) (Supplementary Figure 2A), using K-nearest neighbour imputation with three neighbours. Data imputation was applied to 39 variable including demographic data (age and sex), cognitive tests used to calculate general cognitive ability score (GCAS) and imaging derived phenotypes representing structure and functional front-parietal connectivity (FPC IDPs). Subsequently, the same analyses as with complete cases, were performed using imputed data, and are reported in Supplementary Materials. Further information, all codes and figures are available from https://github.com/InFlamUOB/CognitiveReserve.

### General cognitive ability score (GCAS)

To capture heterogeneity in cognitive functioning in the UK Biobank ageing cohort, we calculated and employed a general cognitive ability score (GCAS), an approach previously suggested for studies considering shared genetic, environmental and sociodemographic influences (Fawns-Ritchie and Deary, 2020; Lyall et al., 2016). These was done using principal component analysis (PCA), saving scores based on the first principal component, which accounted for 30.9% of the variance, similarly to previous reports (Fawns-Ritchie and Deary, 2020; Lyall et al., 2016). Eigenvalues and scree plots can be found in Supplementary Figure 4 and Supplementary Table 2. The calculated GCAS represents composite cognitive ability and as such was subsequently used as a measure of cognitive outcomes.

### GWAS derived SNPs

The genetic analyses were carried out based on genotype data derived from blood samples collected during baseline visit, in total UK Biobank includes genetic data for 488,377 participants. In accordance with UK Biobank protocols, following DNA extraction, the genotyping was performed using two different arrays, the UK BiLEVE Axiom array and the UK Biobank Axiom array. The full details of the genotyping, quality control and imputation have been previously published (Bycroft et al., 2018) and are available from the UK Biobank showcase (https://biobank.ndph.ox.ac.uk/ukb). In order to extract genetic features, we selected specific genetic variants which have been implicated in noradrenergic and dopaminergic neurotransmission. The final gene set, total of 154 genes (464,939 SNPs) was selected based on search performed using the Molecular Signature Database (MSigDB; https://www.gsea-msigdb.org/gsea/msigdb/), and published literature (Bralten et al., 2013; van Donkelaar et al., 2020) and can be found in in https://github.com/InFlamUOB/CognitiveReserve. In the analysis, we have included genes associated with both noradrenergic and dopaminergic genetic pathways as the two neurotransmitters share both signalling and metabolic pathways due to overlapping biosynthesis and the known biochemical similarity of some of the transporters and receptors (Carboni et al., 1990; Cornil and Ball, 2008; Wedemeyer et al., 2007). Thus, separating the two genetic pathways is highly problematic, if not impossible. SNPs within all identified genes and their flanking regions (to include regulatory sequences) were selected for GWAS. These SNPs were then tested for associations with the previously mentioned metrics: cognitively enriched environments (education), cognitive outcomes (measures of performance on 10 cognitive tests) and fronto-parietal connectivity (structural and functional connectivity IDPs), using genome-wide association analysis (GWAS). More specifically the tools PLINK (Purcell et al., 2007) for pre-processing and quality control and REGENIE (Mbatchou et al., 2021) for GWAS were used. REGENIE is a computationally efficient machine learning method that reduces the genotype space into local blocks (of 1000 variants) before performing association testing. After GWAS, chosen SNPs were filtered (LOG10P > 4.5), yielding 53 SNPs. Summary statistics of selected SNPs can be seen in https://github.com/InFlamUOB/CognitiveReserve. In addition, to control for potential confounding effects in our analysis, we included SNPs e3e3, e3e4 and e4e4 from apolipoprotein-E (APOE), which is known to be a risk factor for dementia (Heffernan et al., 2016). The distribution of all final variables introduced in the model encompassing (1) GWAS derived SNPs (2) demographics (age and sex) (3) general cognitive ability score (GCAS) from cognitive tests (4) imaging derived phenotypes representing structure and functional front-parietal connectivity (FPC IDPs) and (5) cognitive reserve information using education as proxy can be seen in Supplementary Figure 5.

### Statistical analysis

All analyses were performed in R version 4.2.0

### Modelling

As our objective is to model interactions between neurobiological features (both genetic and connectivity measures) and education (proxy of CR) on cognitive outcomes, we built regression models using general cognitive ability score (GCAS) as outcome. Using *workflowsets* and *tidymodels*, 5 different algorithms (generalized linear model (*glm* package), LASSO (Tibshirani, 1996) (*glmnet* package), a single-hidden-layer neural network (*nnet* package), random forest (*ranger* package) and xgboost (*xgboost* package), were fitted to 14 different combinations of the features above extracted. More precisely, 5 different algorithms were run on data comprising: SNPs, FPC IDPs, AgeSex, SNPs x FPC IDPs, SNPs x CR (education), SNPs x AgeSex, FPC IDPs x CR (education), AgeSex x CR (education), SNPs x FPC IDPs x CR (education), SNPs x FPC IDPs x AgeSex, SNPs x AgeSex x CR (education), FPC IDPs x AgeSex x CR (education as proxy), SNPs x FPC IDPs x AgeSex x CR (education) information. In each algorithm and feature combination dataset (69 in total) (Figure 1), hyperparameters were tuned through *grid.search* and 100 cross-validation resamples fitted per model. To cut down on time, *tune_race_anova* (Kuhn, 2014) was used which eliminates tuning parameter combinations that are unlikely to be the best results. To do so*, tune_race_anova* uses a repeated measure ANOVA model just after an initial number of resamples have been evaluated. Performance was evaluated through Mean Absolute Error (MAE), where the sum of absolute errors between predicted GCAS and true GCAS is divided by the sample size.

**Figure 1.**
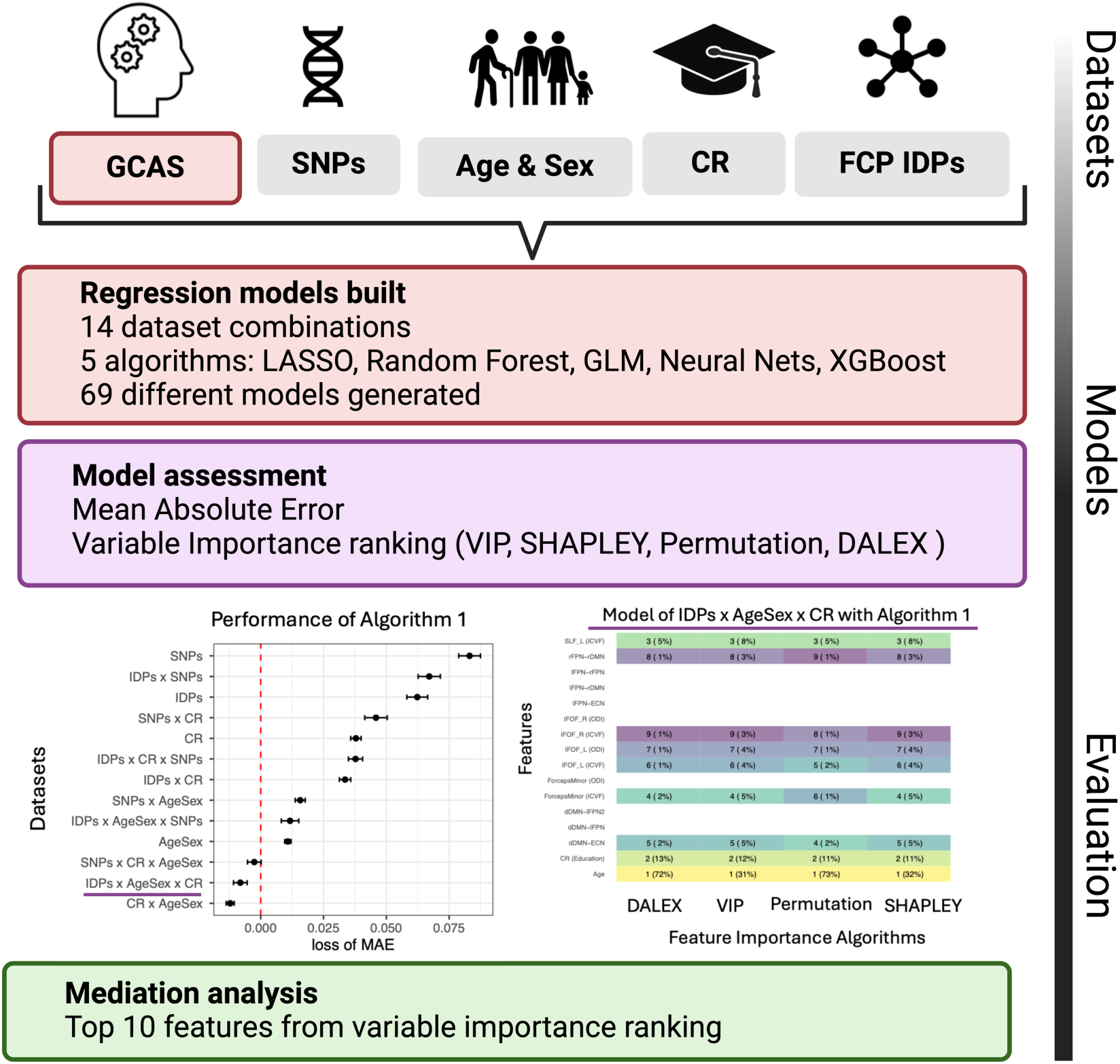
Study Design. We explored the interplay neurobiological features (genetic variants in noradrenergic signalling, structural and functional fronto-parietal connectivity) and education (proxy of CR) by testing regression models predicting general cognitive ability score (GCAS) as outcome. By splitting features into 4 datasets: GWAS derived SNPs, AgeXSex, CR with Education as proxy and imaging derived phenotypes representing structure and functional front-parietal connectivity (FPC IDPs), we generated 14 different dataset combinations. These 14 different feature combinations were then modelled with 5 different algorithms (Least Absolute Shrinkage and Selection Operator (LASSO), Generalized Linear Models, XGBoost, Random Forest and Neural Nets), generating 69 different models. Behaviour of these models was then evaluated through performance, using mean absolute error and feature importance. Given the disparity in modelling algorithms, different feature importance algorithms were also used (DALEX, model dependent variable importance assessment, permutation and SHAPLEY), and results were summarised as ranking scores. Top 10 features (sex and 9 neurobiological variables) selected as most important (top 2) per variable importance method, algorithm and dataset were then studied as mediators (9 neurobiological variables only) between education (CR) and GCAS and including age as covariate.

### Variable importance

To better understand the behaviour of the models, variable importance of each of the 69 different algorithm and variable combinations was assessed using 4 different feature importance methods (Elhabr, 2020). These include model-agnostic algorithms such as the explainable AI package *DALEX* (Biecek, 2018), shapley values (Sundararajan and Najmi, 2020) and permutation related variable importance or model-specific algorithms such as model related variable importance. The loss function in variable importance computations for the model-agnostic methods was minimization of the root mean-squared error (RMSE). The top 9 ranking variables for each of the variable importance assessments in each algorithm and dataset was then visualized in a heatmap, and next the 10 variables ranked the highest (most amount of times appearing in top 2) were selected. These included Sex and 9 neurobiological variables. The 9 variables theoretically relevant to the tested Robertson’s theory (SNPs representing genetic variants in noradrenergic signalling, measures of structural and functional frontoparietal connectivity) were followed through for mediation analysis.

### Mediation

Finally, to assess a possible relationship between education (used here as proxy of CR) on cognitive outcomes, we tested those features that came up as highly important in our models through mediation analysis (VanderWeele, 2016). Mediation analysis was used to assess the magnitude of the data-driven pathways and mechanisms selected as relevant in a data-driven way and how they may affect general cognitive ability score and education relationship. Mediation was performed using libraries *psych* and *mediation* where all 9 selected, consistently important variables for GCAS prediction (as described above, most frequent top 2 important variables in each of the variable assessment models across all datasets and algorithms) were studied. Given age appeared as the most important variable across all models where it was included, we included age as a covariate in our mediation analysis (partialling out its contribution) in our mediation analysis. The mediation diagrams for all variables can be seen in https://github.com/InFlamUOB/Cognitive Reserve with the mediation effect bootstrapped 10,000 times.

## RESULTS

### Pre-processing

From an initial cohort of 502,505 participants, 48,561 with imaging data were identified for the purpose of the current study. From these, 26,227 had complete education information. Missing values assessment can be seen in (Supplementary Figure 1 and 2), with remaining missing values (20.2%) both imputed using k-nearest neighbours (Supplementary Figure 3) and deleted (complete-case analysis) as reported below. Taking advantage of genotypic data, SNPs related to noradrenergic and dopaminergic neurotransmission were extracted as well as APOE alleles e3e3, e3e4 and e4e4 given their close relationship to cognitive decline and dementia (Heffernan et al., 2016; Veldsman et al., 2020). In total 12,076 participants with 25 features associated to brain imaging data, 56 genetic data, 2 demographics and education were selected with their distribution available in Supplementary Figure 4.

### Modelling

To understand what data better captured general cognitive ability, different models were built predicting GCAS using both different feature combinations and modelling strategies (Figure 1). More specifically, generating all different feature combinations between brain imaging, genetic, demographics and education yielded 14 different datasets as seen in Methods. All these datasets were then trained using 4 distinct machine learning algorithms combining both non-linear and linear methods (Random Forest, Neural Nets, LASSO and GLM), creating 69 different models. After splitting our dataset into training (5/6) and testing (1/6) data, models were trained, with hyperparameter tuning and resampling (100 fold-cross validation).

Final performance was assessed in the test set (Figure 2) and feature importance through 4 metrics (vip, permutation, shapley and DALEX) for all models is reported (Figure 2).

**Figure 2.**
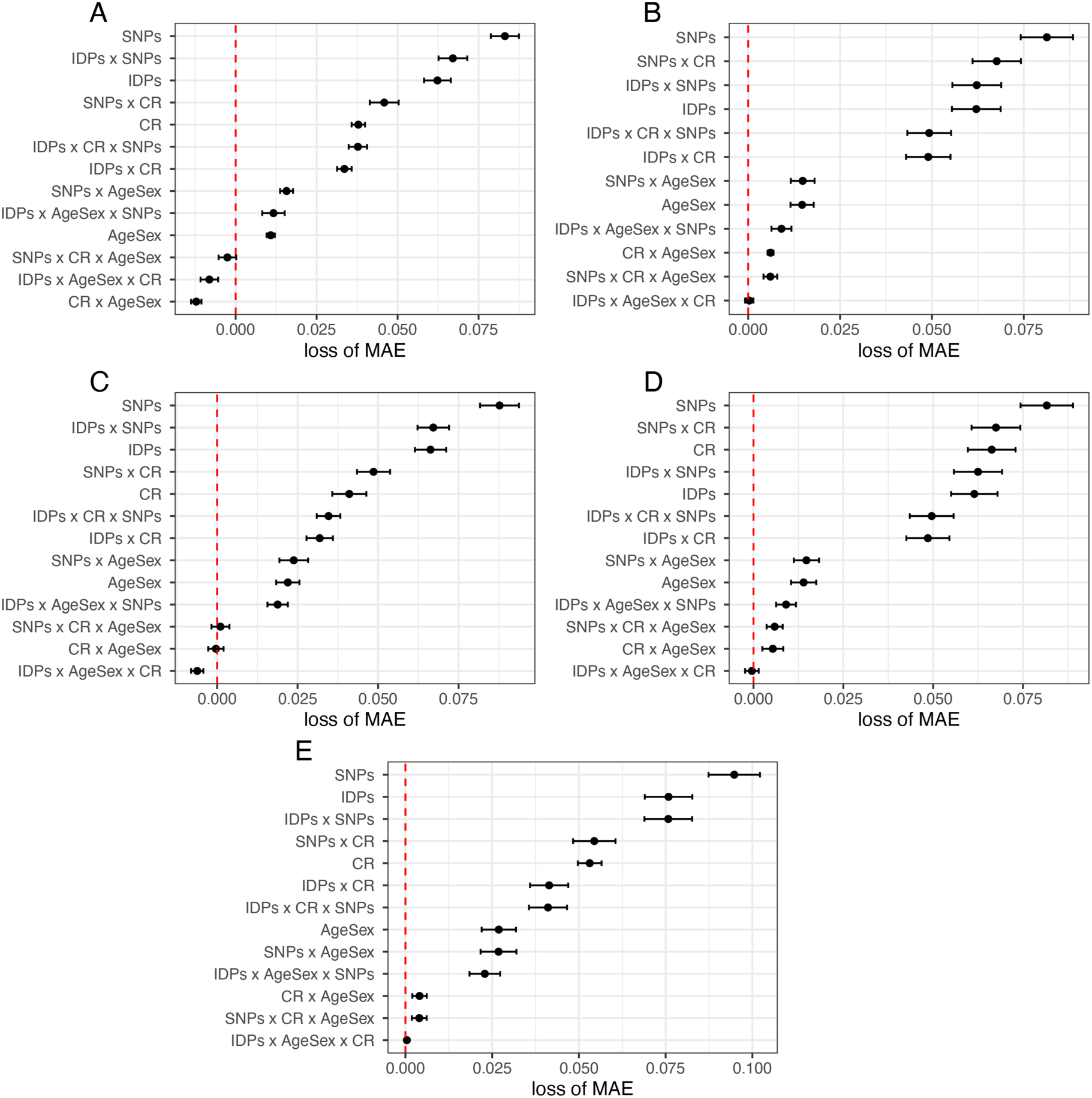
Results from All Regression Models. Mean Absolute Error (MAE) of model combinations of AgeXSex, front-parietal connectivity imaging derived phenotypes (FPC IDPs) and SNPs data for each of the different algorithms (A. Neural Nets, B. LASSO, C. Random Forest, D. GLM and E. XGBoost), using the model with all datasets included as reference value (red line). Error bars correspond to the 100 fold cross-validated performance results. In total, 69 models were analysed (with unique feature and algorithm combination). As a general trend, SNPs possess the least prediction ability, with the highest mean average error across all algorithms (higher loss of MAE with respect to baseline 0 (which is all datasets together). In contrast, including AgeXSex x CR (education) guarantees a better performance across all datasets, with IDPs x AgeSex x CR (education) generally yielding the best performance. All hyperparamters were tuned using tune race_anova with best performing hyperparameter combinations used and test set performance reported here.

A combination of imaging, demographic and education data was seen to be the best performing model in all algorithms, with these feature combinations yielding better performance than combining all features together (reference line in Figure 2). To better understand feature importance per model and algorithm, heatmaps were used to visualize results, with information on the models generated form imaging, demographic and education data seen in Figure 3 as an example. In this heatmap, the ranking of the top 9 features per feature importance methods applied to each model is reported, with Age and Education scoring first and second in all algorithms and methods but less consensus seen in the ranking of imaging data, with varying agreement between algorithms and feature importance methods. Results from the other models can be found in: https://github.com/InFlamUOB/CognitiveReserve

**Figure 3.**
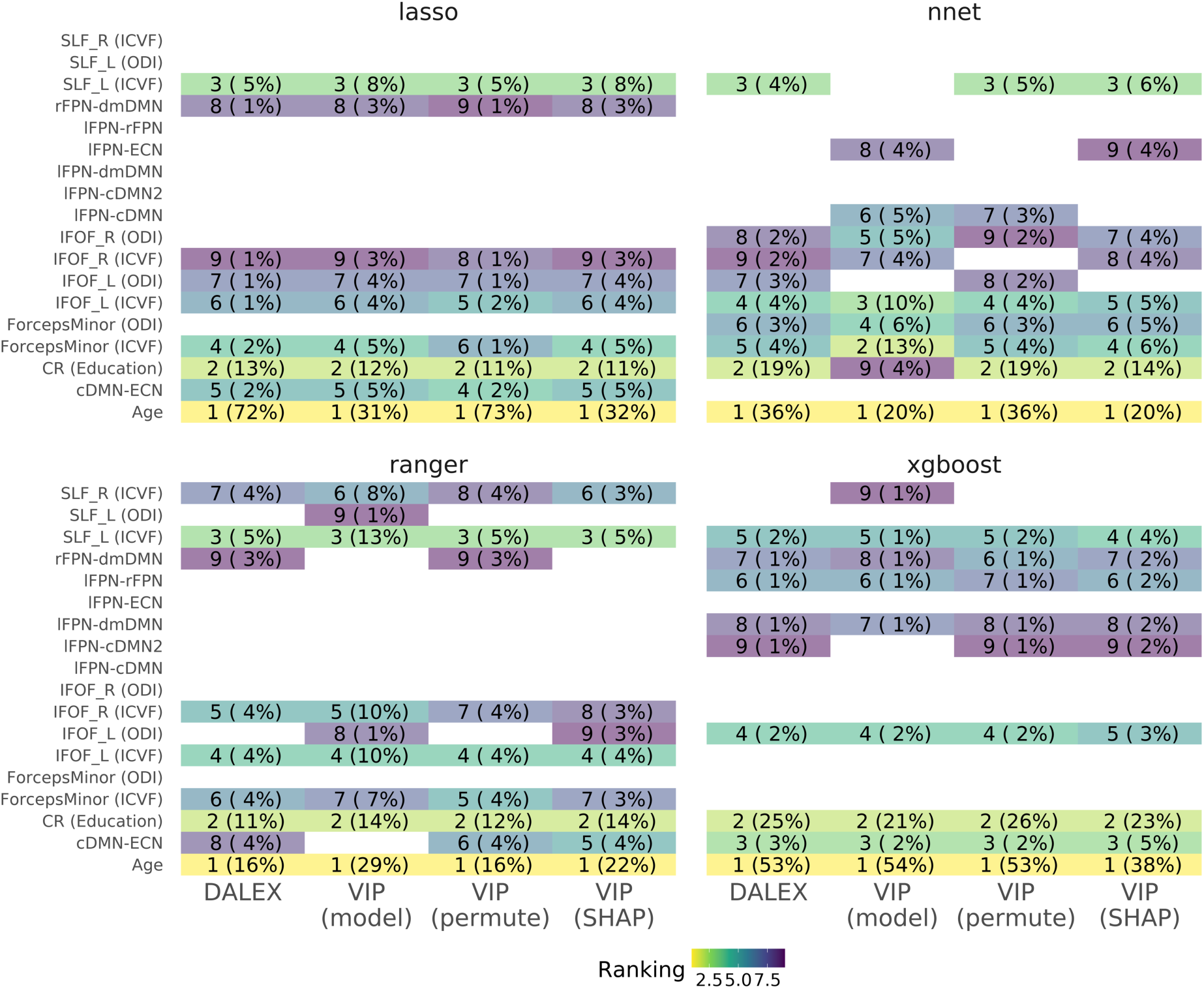
Feature Importance Testing. Heatmap of feature importance for the 4 out of the 5 different algorithms (LASSO and GLM as both linear models, are here only represented as LASSO results) and data comprising features from AgeXSex, CR with education as proxy and front-parietal connectivity imaging derived phenotypes (FPC IDPs). Most important features are ranked using 4 different feature importance algorithms (DALEX, model specific variable importance (VIP), permutation and Shapley values), with most important being 1 (yellow) and least 9 (purple). Only the 9 most important features for each models are here reported. All model-agnostic methods (permutation, DALEX and Shapley) had variable importance assessed though root mean-squared error (RMSE) and the absolute values of importance were assessed as ranking. In brackets, the normalized absolute importance value is reported as percentage.

### Mediation

Finally, wanting to better understand the relationship between education and cognitive outcome, we proceeded to explore the most frequent important features extracted from each variable importance method for each algorithm and dataset in a mediation model. Age was found as the most important variable across models and datasets when included (Figure 3). Thus, taking into account relevance of age to studied hypothesis and employed study sample (ageing UK Biobank cohort including middle age and older adults), we include age in our mediation analysis as an extra covariate and evaluate the top 9 frequent neurobiological features as mediators. More specifically, the direct effects of education (CR proxy) on general cognitive ability score partialling out the effect of each of these features univariately was evaluated with the main effects reported in Table 1. Following that seen as best model (FPC IDPs, education and Age features), the variable seen to have a significant mediation effect was rFPN-dmDMN connectivity (Figure 4, Table 1). The results based on the imputed data are reported in the Supplementary Figure 6. rFPN-dmDMN connectivity was found as a significant mediator in both analyses i.e., using only complete cases and imputed data, thus ascertaining the robustness of findings.

**Figure 4.**
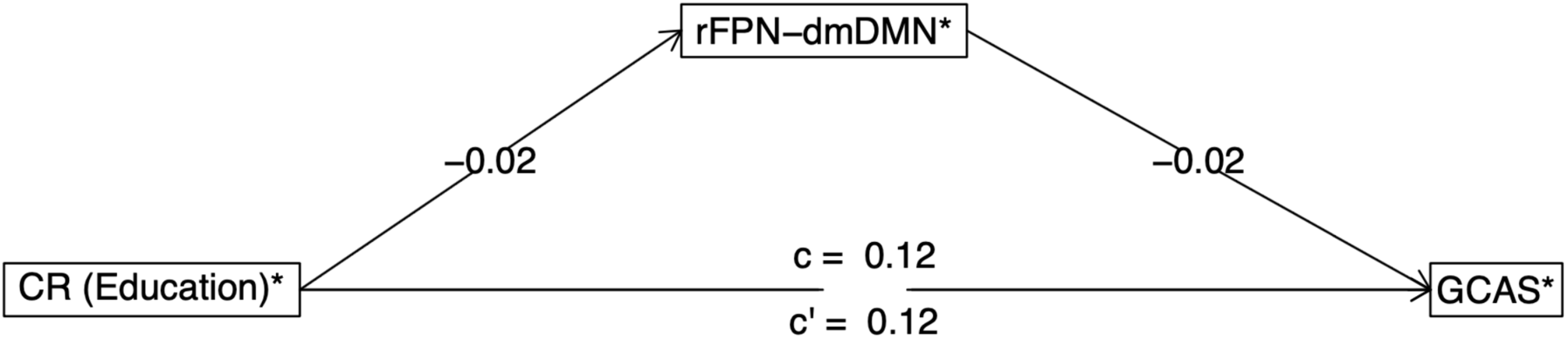
Mediation Analysis Results. rFPN-dmDMN connectivity was found to be a significant mediator with education (proxy of CR) as predictor of General Cognitive Ability Score (GCAS) when controlling for Age (95% CI not crossing 0). Mediation diagram as extracted from mediate.diagram function from the psych package. Significant mediator (95% CI not crossing 0) when controlling for Age.

**Table 1.** Mediation results using *psych* package. Top 9 most frequent important variables (top 2 in ranking) were included in a model as mediators with education (CR) as predictor, General Cognitive Ability Score (GCAS) as outcome and Age as covariate. Bootstrapped confidence intervals for the indirect effect reported as lower CI (2.5%) and upper (97.5%). Significant mediators (95% CI not crossing 0).

| Mediator | Effect of CR on mediator (a) | Effect of mediator (b) | Indirect effect (ab) | 95% CI |  |
| --- | --- | --- | --- | --- | --- |
|  |  |  |  | Lower | Upper |
| SLF_L (ICVF) | 0.0039 (0.006) | 0.0436 (0.0057) | 2e-04 (3e-04) | -3e-04 | 7e-04 |
| rs10224002(PRKAG2) | -0.0014 (0.0016) | 0.0193 (0.021) | 0 (1e-04) | -2e-04 | 1e-04 |
| rs2888788(PRKAG2) | -0.0058 (0.0031) | -0.0385 (0.011) | 2e-04 (1e-04) | 0 | 5e-04 |
| rs10254101(PRKAG2) | 0.0025 (0.003) | 0.0251 (0.0115) | 1e-04 (1e-04) | -1e-04 | 3e-04 |
| IFOF_L (ICVF) | 0.0058 (0.006) | 0.0281 (0.0057) | 2e-04 (2e-04) | -2e-04 | 5e-04 |
| rFPN-dmDMN | -0.018 (0.0062) | -0.0202 (0.0055) | 4e-04 (2e-04) | 1e-04 | 7e-04 |
| rs1872691(BRD7-ADCY7) | 0.003 (0.0027) | 3e-04 (0.0124) | 0 (1e-04) | -1e-04 | 1e-04 |
| IFOF_R (ICVF) | -0.0031 (0.0059) | 0.0286 (0.0057) | -1e-04 (2e-04) | -5e-04 | 3e-04 |
| rs3813755(ADCY7) | 0.0029 (0.0027) | 0.0046 (0.0125) | 0 (1e-04) | -1e-04 | 1e-04 |

### Code Availability

All code and figures can be found in: https://github.com/InFlamUOB/CognitiveReserve.

## DISCUSSION

Using robust machine learning methods and mediation analysis applied to data from the UK Biobank Brain Imaging study (Miller et al., 2016), we tested here combinatorial effects of neurobiological features (genetic variants in noradrenergic signalling, measures of structural and functional frontoparietal connectivity) and education (used here as proxy of CR) on cognitive outcomes in the UK Biobank ageing cohort. Initially, several different models were built with a large number of features. We then selected a small number of features, related to genetic variance, fronto-parietal connectivity alongside sex and age, for subsequent analysis using mediation testing. A variety of machine learning algorithms and feature importance methods were used given that there is no algorithm able to fit all data perfectly (“No Free Lunch Theorem”(Belkin et al., 2019)). Therefore, testing more than one algorithm and more than one approach to interpret feature importance provides us with the possibility of acknowledging many more possible underlying relationships. Also feature importance was especially relevant for black box algorithms such as neural nets and random forest where behaviour of the model is more difficult to grasp. Based on these analyses we show that (1) interactions between education and neurobiological variables more fully explain general cognitive performance on multiple tests (general cognitive ability score) than either factor alone, and (2) among the examined neurobiological features measures of functional fronto-parietal connectivity are the strongest mediators of the effect of education on cognitive outcomes.

Our main findings point to fronto-parietal resting state functional connectivity measures (rFPN-dmDMN, Figure 4) representing connections between the default mode network and the right fronto-parietal network as significant mediator between education and cognitive outcomes in UK Biobank ageing population sample. This suggests that functional connectivity between several neural networks, including right fronto-parietal network, might be sensitive to the effects of education (proxy of CR) on cognition in middle and old age. Thus, alongside our previous studies (Shalev et al., 2020; Brosnan et al., 2022), we provide here evidence in support of Robertson’s theory of CR (Robertson 2013, 2014). Robertson’s theory stipulates that at the neural level the lifelong exposure to cognitive stimulation strengthens the right hemisphere fronto-parietal networks, which in turn offsets the symptoms of cognitive ageing. Our previous work (Brosnan et al., 2022; Shalev et al., 2020) and that of others (van Loenhoud et al. 2017), indeed point to the right hemispheric fronto-parietal networks as neural substrates of CR. By contrast here we linked the phenomenon of CR to not only the right fronto-parietal network but also the default mode network encompassing additional bilateral fronto-parietal regions, as well as the connectivity between these two networks. The Robertson’s theory proposes that exposure to enriched environments acts via core cognitive processes, including alertness, sustained attention, and awareness which indeed are known to be subserved by the right fronto-parietal networks (e.g., Beck et al., 2001; Naghavi and Nyberg, 2005; Hester et al., 2005; Singh-Curry and Husain, 2009; Stuss, 2011). However, the default mode networks implicated by our current study is also functionally linked to the cognitive processes described by Robertson’s theory. Furthermore, similarly to fronto-parietal network, the default mode network is known to be modulated by the locus coeruleus-noradrenergic (LC-NE) system (Bär et al., 2016; Minzenberg et al., 2018; Robbins & Arnsten, 2009; Suttkus et al., 2021), which is pivotal to Robertson’s theory.

It should be also noted that while the mentioned above studies, supporting Robertson’s theory, only examined the beneficial effects of CR on structural measures within fronto-parietal networks (van Loenhoud et al. 2017; Shalev et al., 2020; Brosnan et al., 2022;), the current study links the phenomenon of CR (measured here by proxy of education) to measures of functional connectivity. Several previous studies in non-demented older adults linked measures of functional connectivity to offsetting effects of cognitive reserve either on global cognition or memory performance (e.g., Varela-Lopez et al., 2022; Boyle et al., 2023; Franzmeier et al., 2018; Fleck et al., 2019). However, most of these studies either explored connectivity measures derived from a task-based fMRI and/or the analysis were primarily restricted to measures of within-network connectivity, and additionally the previous findings were limited by relatively small sample sizes (less than 100 participants). By contrast we explored here between networks connectivity in a large cross-sectional UK Biobank cohort. But what is the most striking about our findings is that they point to the default mode network as the key player in between networks interactions underlying cognitive reserve. As noted above this expands the Robertson’s proposal (Robertson, 2014), which specifically stipulates that the hypothetical CR network consists of interconnected right lateral prefrontal lobe and right inferior parietal lobe regions. Our findings expand the CR network, to include bilaterally dorsal medial prefrontal cortex and posterior cingulate cortex.

In the context of ageing, the default mode network is perhaps the most studied of the resting state networks (Jiang et al., 2020; Staffaroni et al., 2018) with functional changes linked to cognitive decline and dementia (Vidal-Peneiro et al., 2014; Andrews-Hanna et al., 2007; Damoiseaux et al., 2008; Bluhm et al., 2008; Mevel et al., 2011; Hafkemeijer et al., 2012; Grieder et al., 2018). In addition, when examining a small sample of Alzheimer’s patients, Bozzali and colleagues (2015) previously found evidence that cognitive reserve modulate connectivity within the default mode network. Finally, our results expand on newly published findings linking changes in connectivity between the default mode network and frontoparietal networks to cognitive decline in elderly (Koshino et al., 2023) as well as to the effects of beta-amyloid on cognitive status in Alzheimer’s patients (Zhukovsky et al., 2023) in line with classical account of cognitive reserve (Stern et al., 1992; Stern, 2012) i.e., mitigating accumulated neuropathology.

To our knowledge, only one previous study examined underpinnings of the effects of CR on cognitive outcomes in the UK Biobank cohort (Jin et al., 2023). However, the scope, research objectives and methods employed by Jin et al (2023) and here are very different. Jin and colleagues specifically conducted a series of studies systematically exploring the effects of different CR proxies (education, leisure activities, fluid intelligence, social interactions, and physical activity) on the relationship between structural brain measures (global volume, regional volumetric measures and cortical thickness within brain areas know to deteriorate in dementia patients) and cognitive outcomes indexed by performance on tests within 4 separate cognitive domains. The main aim of their research was to explore the reported inconsistencies in the literature concerning the moderating effects of various CR proxies on the links between brain structure and cognitive abilities in an ageing population. This was motivated by the premise that CR explains the discrepancy between accumulated brain damage and observed cognitive performance in accordance with classical theory of cognitive reserve as proposed by Stern and based on his early work in Alzheimer’s patients (Stern et al., 1992; Stern, 2012). By contrast our analyses were motivated by the Robertson’s proposal (2013, 2014) that CR, here represented by education strengthen the right fronto-parietal networks to offset age-related cognitive decline. Thus, the only link between Jin et al (2023) and our study, conducted using data from UK Biobank cohort, is that both provide support for the notion of beneficial effects of CR on cognitive ageing.

One of the objectives of the current study was to explore a premise that an interplay between accumulative effects of CR (represented by education) and genetic variance in noradrenergic signalling on frontoparietal neural networks might offset age-related cognitive decline. While previous studies indeed linked genetic variations in neurotransmitter signalling to heterogeneity in cognitive ageing (Lindenberger et al., 2008; Nyberg et al., 2014; Papenberg et al., 2015a,b), we have not found any evidence of the mediating effects of genetic variability in noradrenergic signalling on interplay between education and cognitive outcomes. It should be noted that the previous evidence comes from hypothesis driven genetic association studies using single-nucleotide polymorphisms (SNPs). By contrast we employed here GWAS based approach to find genetic variants associated with education, cognitive outcomes and fronto-parietal connectivity (structural and functional). While the GWAS analyses were notably restricted to the preselected gene set, 154 genes totalling 464,939 SNPs, the negative findings might not be that surprising considering a relatively small sample comparing to other GWAS studies examining genetic underpinnings of education and cognition (e.g., Okbay et al., 2016; Davies et al., 2016). Thus, the size of study sample constitutes a limitation when exploring genetic influences. The negative findings might also potentially reflect shortcomings of the taken here approach to testing Robertson’s theory. We employed here genetic variance in noradrenergic signalling as variable depicting noradrenergic modulation in Robertson’s theory but it is plausible that the noradrenergic genetic variance does not sufficiently capture cumulative influences of cognitively enriched environments as per Robertson’s theory of cognitive reserve. We have not directly examined here the locus coeruleus function directly indicative of noradrenergic activity. However, it should be noted that the beneficial effects of noradrenergic signalling and the LC function in the context of both cognitive decline and progression of Alzheimer’s disease have been recently questioned by many (for comprehensive review see Mather, 2021). Furthermore, it is plausible that education used as sole proxy of CR does not reflects the complexity of lifelong cognitively stimulating experiences and further work should consider a broader approach to factors contributing to CR. Although, it could be argued that education lessens the decline in cognitive functioning as we age, not only via enhancing neural resources early in life (childhood till early adulthood) with effects lasting later in life, but also at least partially influences professional and lifestyle choices throughout life i.e., other proxy measures of CR. On contrary, some argue that education reflects only individual differences in cognitive abilities that persist from childhood to old age (for review see Cabeza et al., 2018, Lövdén et al., 2020). The assumption is that education attenuates the decline in cognitive functioning as we age, likely via enhancing neural resources early in life (childhood till early adulthood) with effects lasting later in life, Finally, the scope and size of UK Biobank data resources are unprecedented, however similarly to other large scale epidemiological datasets, UK Biobank database includes large proportion of incomplete cases. Here, out of an initial sample of 48,561 participants, only 12,076 complete cases were found (i.e., including all brain imaging data, cognitive, genetic, demographic and education variables). The missing cases cause potential limitations due to loss of statistical power (e.g. in GWAS as discussed above) and by contributing to risk of bias affecting findings. To address that we repeated all analyses using imputed data. Importantly, we again found the measures of fronto-parietal connectivity to be the strongest mediators of the effect of education on cognitive outcomes.

In summary, the data presented here not only add to our understanding of the neurobiological underpinnings of CR but also the role of between network connectivity and the default mode network in cognitive ageing. By employing comprehensive machine learning methods, we have demonstrated that between networks functional fronto-parietal connectivity is a strong mediator of the effect of education used her as proxy of cognitive reserve on cognitive outcomes in ageing population. Our findings provide additional support for Robertson’s theory of CR (Robertson, 2013, 2014) as well as the recent finding in Alzheimer’s patients linking between networks frontoparietal connectivity to offsetting effects against amyloid burden (Zhukovsky et al., 2023). While on one hand the limitation of the current study is that we only examined between network connections, on the other hand several theories of cognitive ageing explicitly address changes in functional correlations between functional networks associated with either preservation or decline in cognitive performance in ageing. Specifically, it has been suggested that processes such as compensation and dedifferentiation both might be accompanied by stronger correlations among functionally unrelated networks or weaker anti-correlations among competing networks (Malagurski et al., 2020a,b; Deery et al., 2023). In this context the default mode network is of particular interest as the existing evidence points to anti-correlations between the default mode network and various other networks including right fronto-parietal networks at rest and during task performance in young healthy participants (Greicius et al., 2003; Fox et al., 2005). Strikingly, it has also been demonstrated that in elderly participants the default mode network is significantly less deactivated during task performance and that this diminished suppression might underly poorer cognitive performance (Damoiseaux et al., 2008; Persson et al., 2007; Ferreira and Busatto, 2013). Thus, future research should address both cross-sectionally and longitudinally whether the offsetting effects of exposure to enriched environments on cognitive functioning in older adults translate into changes in co-activations and anti-correlations between the default mode networks and task-related fronto-parietal networks while participants are performing cognitive tasks using functional MRI.

## DATA AVAILABILITY

This research has been conducted using the UK Biobank Resource (www.ukbiobank.ac.uk). The authors do not have permission to share data. All researchers wishing to use UK Biobank data need to apply for access and gain approval directly from UK Biobank via the Access Management System (AMS). The data access procedures are set up by UK Biobank https://www.ukbiobank.ac.uk/enable-your-research/apply-for-access.

All analysed here data are available from the UK Biobank https://www.ukbiobank.ac.uk/enable-your-research and the corresponding UK Biobank showcase ID fields are listed in the Appendix (Supplementary Materials).

All in house analysis materials, code, figure and summary statistics are available from https://github.com/InFlamUOB/CognitiveReserve.

## ACKNOWLEDGMENTS

This research has been conducted using the UK Biobank Resource under Application Numbers 29447 and 31224.

## SUPPLEMENTARY MATERIALS

**Supplementary Table 1.** Features included in the analysis before preprocessing as separated into the datasets summarized in Figure 1 (Demographics with AgeXSex, imaging derived penotypes representing structure and functional front-parietal connectivity (FPC IDPs), cognitive tests and education information).

| Variables | Total N<br>(complete cases %) | Levels | Total | Missing education information |  |
| --- | --- | --- | --- | --- | --- |
|  |  |  |  | No | Yes |
| Total N (%) |  |  | 48561 | 26227 (54.0) | 22334 (46.0) |
| **Demographics** |  |  |  |  |  |
| Sex | 48561 (100.0) | Female | 25078 (51.6) | 13969 (53.3) | 11109 (49.7) |
|  |  | Male | 23483 (48.4) | 12258 (46.7) | 11225 (50.3) |
| Age | 48561 (100.0) | Mean (SD) | 64.7 (7.7) | 65.1 (7.7) | 64.2 (7.7) |
| **Cognitive Tests** |  |  |  |  |  |
| ReactionTime | 45217 (93.1) | Mean (SD) | 597.1 (110.7) | 605.1 (113.4) | 587.8 (106.8) |
| NumericMemory | 33890 (69.8) | Mean (SD) | 6.7 (1.5) | 6.4 (1.6) | 6.9 (1.4) |
| FluidIntelligence | 44657 (92.0) | Mean (SD) | 6.6 (2.1) | 6.0 (1.9) | 7.2 (2.0) |
| TrailA | 33021 (68.0) | Mean (SD) | 226.3 (90.5) | 231.8 (98.2) | 220.4 (80.9) |
| TrailB | 33021 (68.0) | Mean (SD) | 558.9 (284.0) | 591.2 (313.8) | 524.2 (243.3) |
| MatrixPatterns | 32646 (67.2) | Mean (SD) | 7.9 (2.1) | 7.4 (2.1) | 8.5 (2.0) |
| TowerRearr | 32363 (66.6) | Mean (SD) | 9.8 (3.2) | 9.5 (3.3) | 10.2 (3.1) |
| SymbolSubs | 32668 (67.3) | Mean (SD) | 18.7 (5.3) | 18.0 (5.4) | 19.5 (5.2) |
| PairedAssoc | 33021 (68.0) | Mean (SD) | 6.8 (2.6) | 6.4 (2.7) | 7.4 (2.5) |
| PairMatch | 45523 (93.7) | Mean (SD) | 3.7 (3.0) | 3.7 (3.0) | 3.6 (3.0) |
| **FPC IDPs** |  |  |  |  |  |
| ForcepsMinor (ICVF) | 37336 (76.9) | Mean (SD) | 0.5 (0.0) | 0.5 (0.0) | 0.5 (0.0) |
| IFOF_L (ICVF) | 37336 (76.9) | Mean (SD) | 0.5 (0.0) | 0.5 (0.0) | 0.5 (0.0) |
| IFOF_R (ICVF) | 37336 (76.9) | Mean (SD) | 0.5 (0.0) | 0.5 (0.0) | 0.5 (0.0) |
| SLF_L (ICVF) | 37336 (76.9) | Mean (SD) | 0.6 (0.0) | 0.6 (0.0) | 0.6 (0.0) |
| SLF_R (ICVF) | 37336 (76.9) | Mean (SD) | 0.6 (0.0) | 0.6 (0.0) | 0.6 (0.0) |
| ForcepsMinor (ODI) | 37336 (76.9) | Mean (SD) | 0.2 (0.0) | 0.2 (0.0) | 0.2 (0.0) |
| IFOF_L (ODI) | 37336 (76.9) | Mean (SD) | 0.2 (0.0) | 0.2 (0.0) | 0.2 (0.0) |
| IFOF_R (ODI) | 37336 (76.9) | Mean (SD) | 0.2 (0.0) | 0.2 (0.0) | 0.2 (0.0) |
| SLF_L (ODI) | 37336 (76.9) | Mean (SD) | 0.2 (0.0) | 0.2 (0.0) | 0.2 (0.0) |
| SLF_R (ODI) | 37336 (76.9) | Mean (SD) | 0.2 (0.0) | 0.2 (0.0) | 0.2 (0.0) |
| cDMN-rFPN | 40012 (82.4) | Mean (SD) | -0.8 (1.0) | -0.7 (1.0) | -0.8 (1.0) |
| dDMN-IFPN | 40012 (82.4) | Mean (SD) | -0.5 (0.9) | -0.5 (0.9) | -0.5 (0.9) |
| cFPN-IFPN | 40012 (82.4) | Mean (SD) | 1.3 (1.0) | 1.3 (1.0) | 1.2 (1.0) |
| cDMN-mtDMN | 40012 (82.4) | Mean (SD) | 1.6 (0.9) | 1.6 (0.9) | 1.5 (0.9) |
| rFPN-mtDMN | 40012 (82.4) | Mean (SD) | 0.3 (0.9) | 0.3 (0.9) | 0.3 (0.9) |
| IFPN-mtDMN | 40012 (82.4) | Mean (SD) | -0.1 (0.9) | -0.2 (0.9) | -0.1 (0.9) |
| cDMN-dmDMN | 40012 (82.4) | Mean (SD) | 2.5 (0.9) | 2.5 (0.9) | 2.5 (0.9) |
| rFPN-dmDMN | 40012 (82.4) | Mean (SD) | 0.3 (1.0) | 0.4 (1.0) | 0.3 (1.0) |
| IFPN-dmDMN | 40012 (82.4) | Mean (SD) | 1.1 (0.9) | 1.1 (0.9) | 1.1 (0.9) |
| mtDMN-dmDMN | 40012 (82.4) | Mean (SD) | -0.8 (0.9) | -0.8 (0.9) | -0.8 (0.9) |
| cDMN-ECN | 40012 (82.4) | Mean (SD) | -1.4 (1.0) | -1.4 (1.0) | -1.5 (1.0) |
| rFPN-ECN | 40012 (82.4) | Mean (SD) | 0.2 (0.9) | 0.2 (0.9) | 0.2 (1.0) |
| IFPN-ECN | 40012 (82.4) | Mean (SD) | -1.2 (1.0) | -1.2 (1.0) | -1.2 (1.0) |
| mtDMN-ECN | 40012 (82.4) | Mean (SD) | -0.0 (1.0) | -0.0 (0.9) | -0.0 (1.0) |
| dmDMN-ECN | 40012 (82.4) | Mean (SD) | 1.4 (0.9) | 1.4 (0.9) | 1.3 (0.9) |

**Supplementary Figure 1A:**
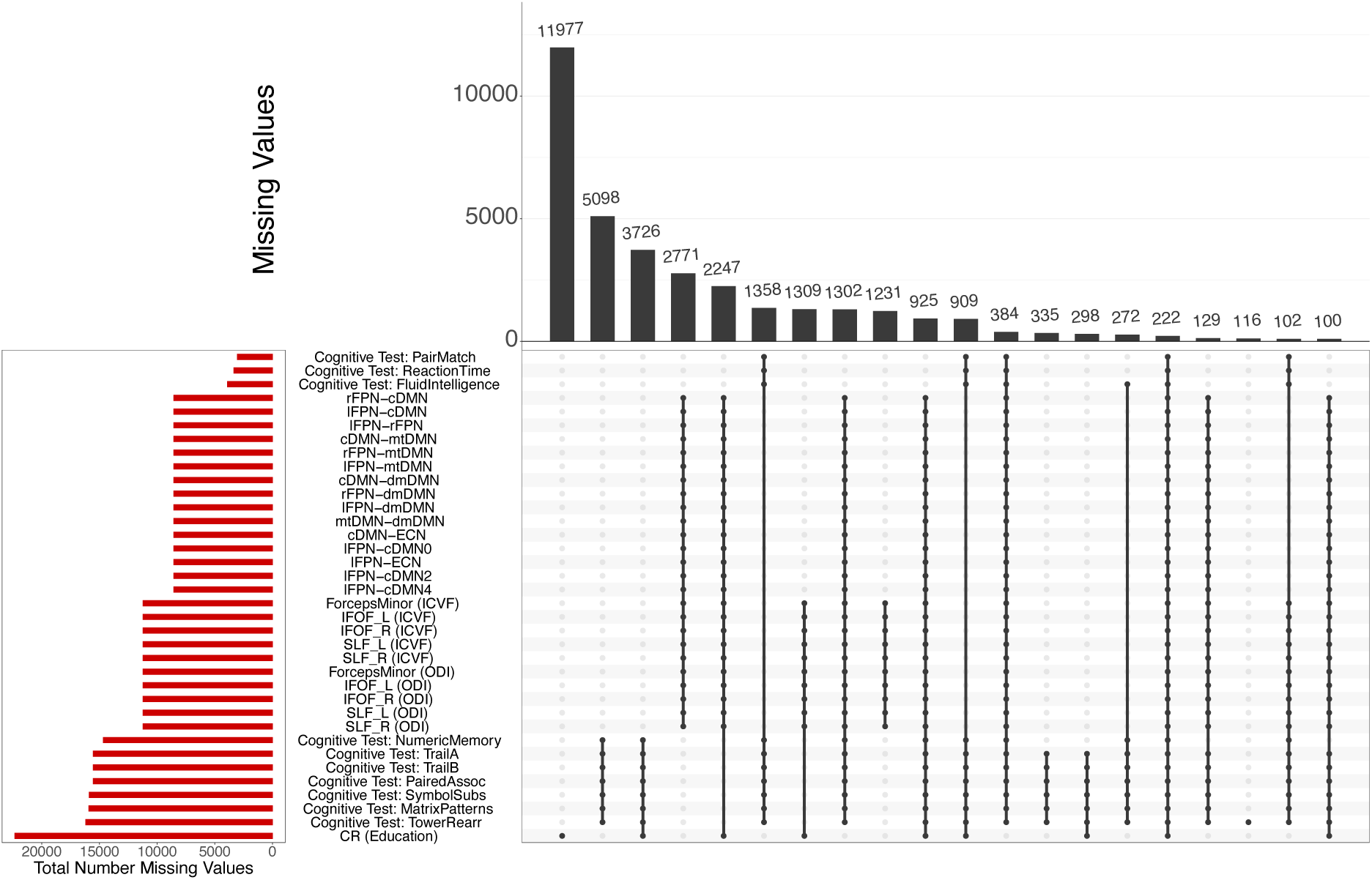
Missing values and missing at random analysis. Missing values pattern exploration using R package *UpSetR* showing the combinations of missingness across cases.

**Supplementary Figure 1B:**
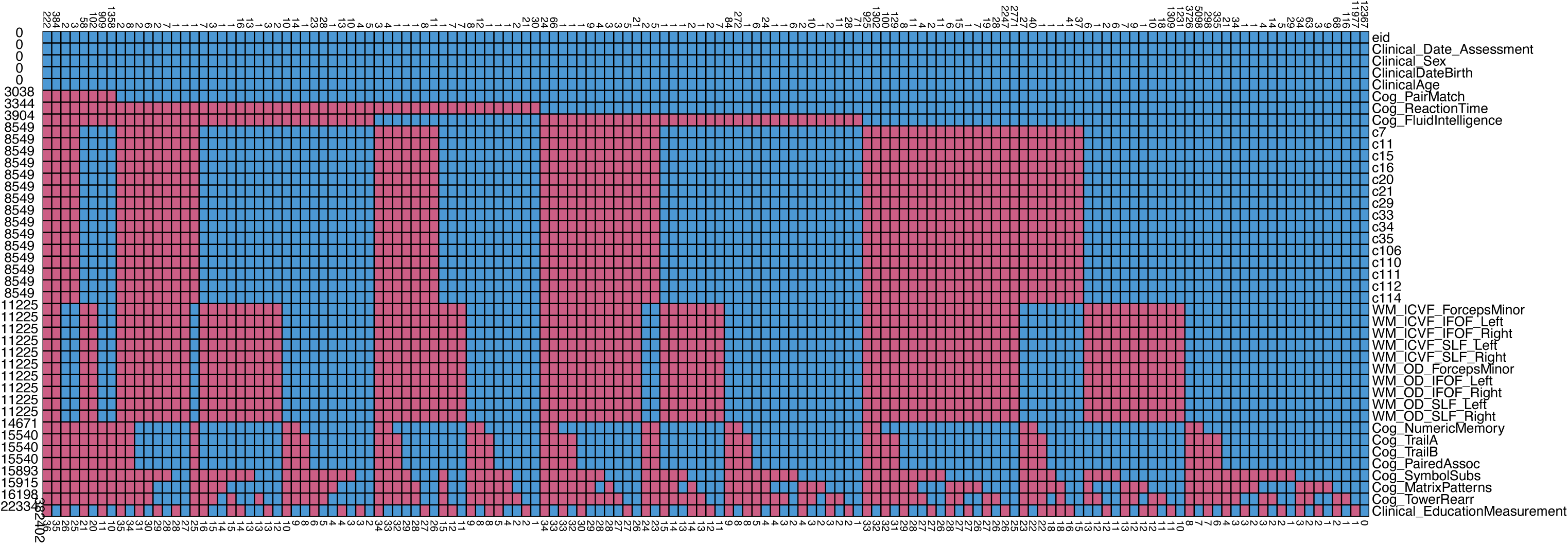
Missing values and missing at random analysis. Missing values pattern exploration using R package *mice* and *finalfit*. Missing values in red and complete in blue, with total numbers of participants found to the left and total missing features (red squares) to the right. Total missing values per feature found at the bottom.

**Supplementary Figure 2.**
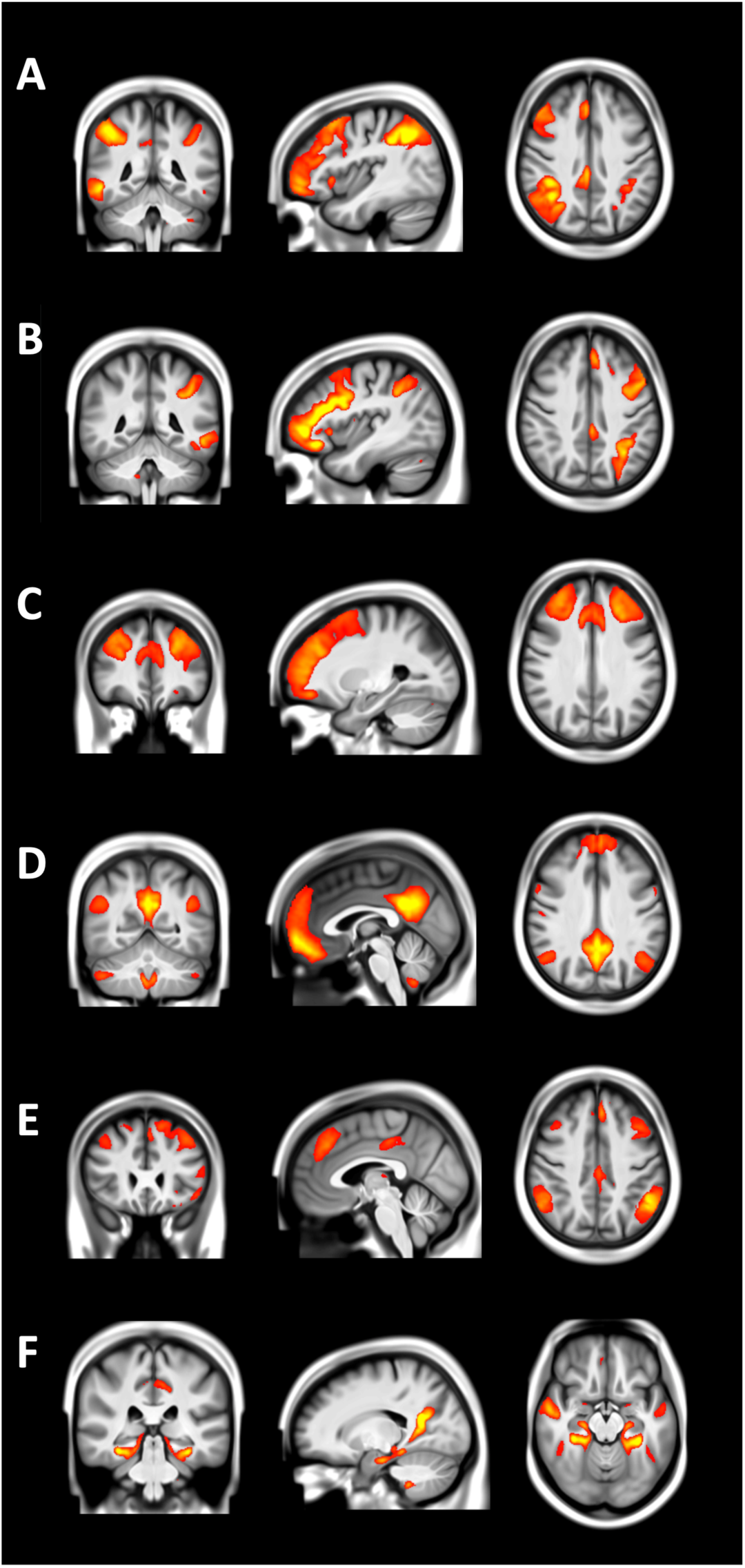
The six ICA components (nodes) representing restig state networks chosen for the purpose of the current study.A. right fronto-parietal network, B. left frontoparietal network, C. executive control network and 3 subsytems of the default mode network: D. core (cDMN), E. dorsomedial prefrontal (dmDMN) and F. medial temporal (mtDMN).

**Supplementary Figure 3:**
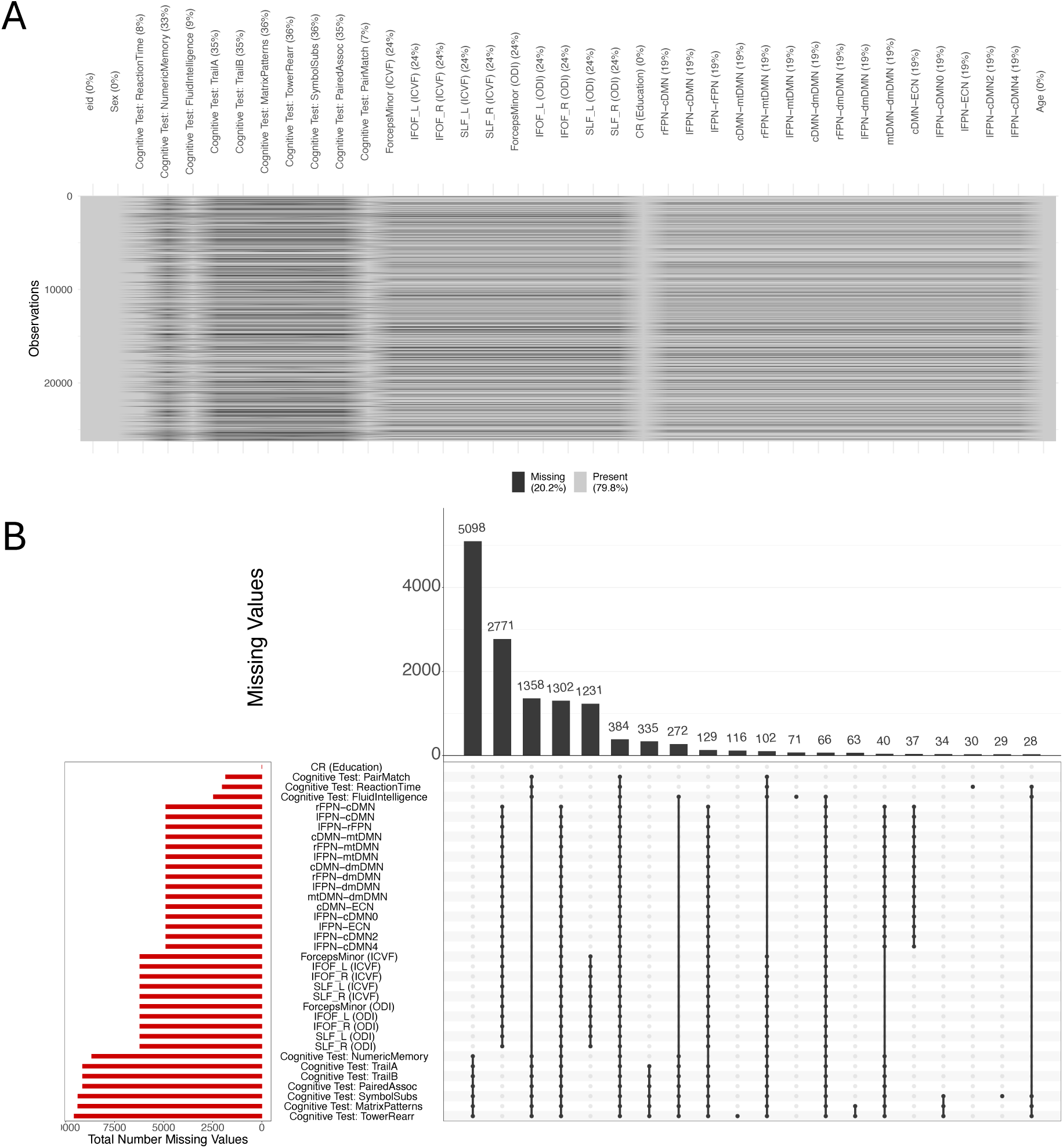
Missing values and missing at random analysis. Missing values pattern exploration using R package *UpSetR* and *naniar*, showing the combinations of missingness across cases. Maximimum of missing cases per column is the cognitive test of tower rearrangement with (36.47%) missing data.

**Supplementary Figure 4.**
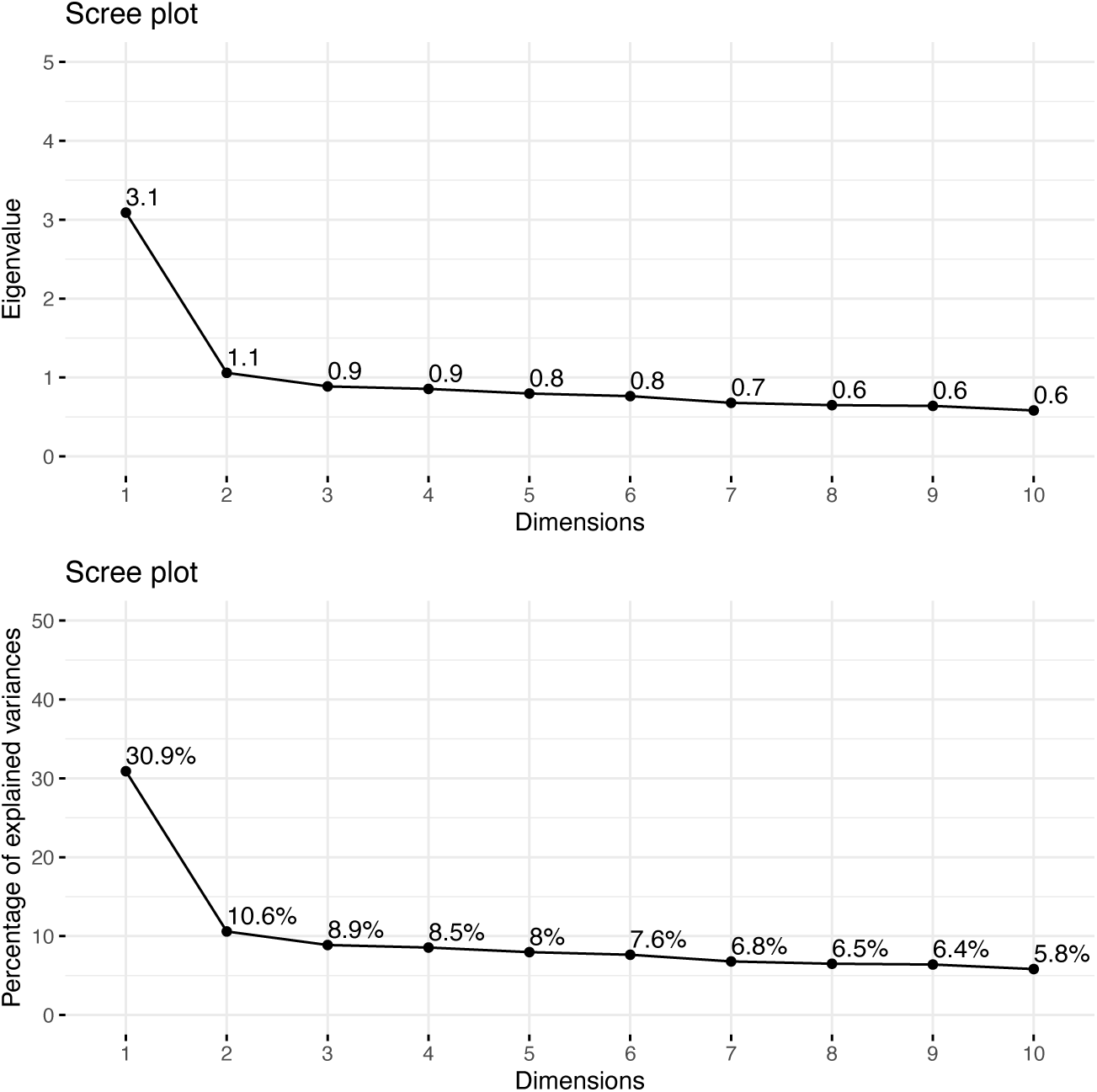
PCA scree plots. See Methods section for further information.

**Supplementary Table 2.** PCA coordinate s(loading* standard deviation) using *factoextra*.

|  | PC 1 | PC 2 | PC 3 |
| --- | --- | --- | --- |
| ReactionTime | -0.384 | 0.496 | -0.473 |
| NumericMemory | 0.512 | 0.429 | 0.084 |
| FluidIntelligence | 0.636 | 0.412 | 0.018 |
| TrailA | -0.544 | 0.379 | -0.069 |
| TrailB | -0.587 | 0.17 | -0.076 |
| MatrixPatterns | 0.646 | 0.185 | 0.01 |
| TowerRearr | 0.626 | 0.002 | -0.077 |
| SymbolSubs | 0.652 | -0.231 | 0.056 |
| PairedAssoc | 0.495 | 0.33 | 0.006 |
| PairMatch | -0.395 | 0.3 | 0.798 |

**Supplementary Figure 5:**
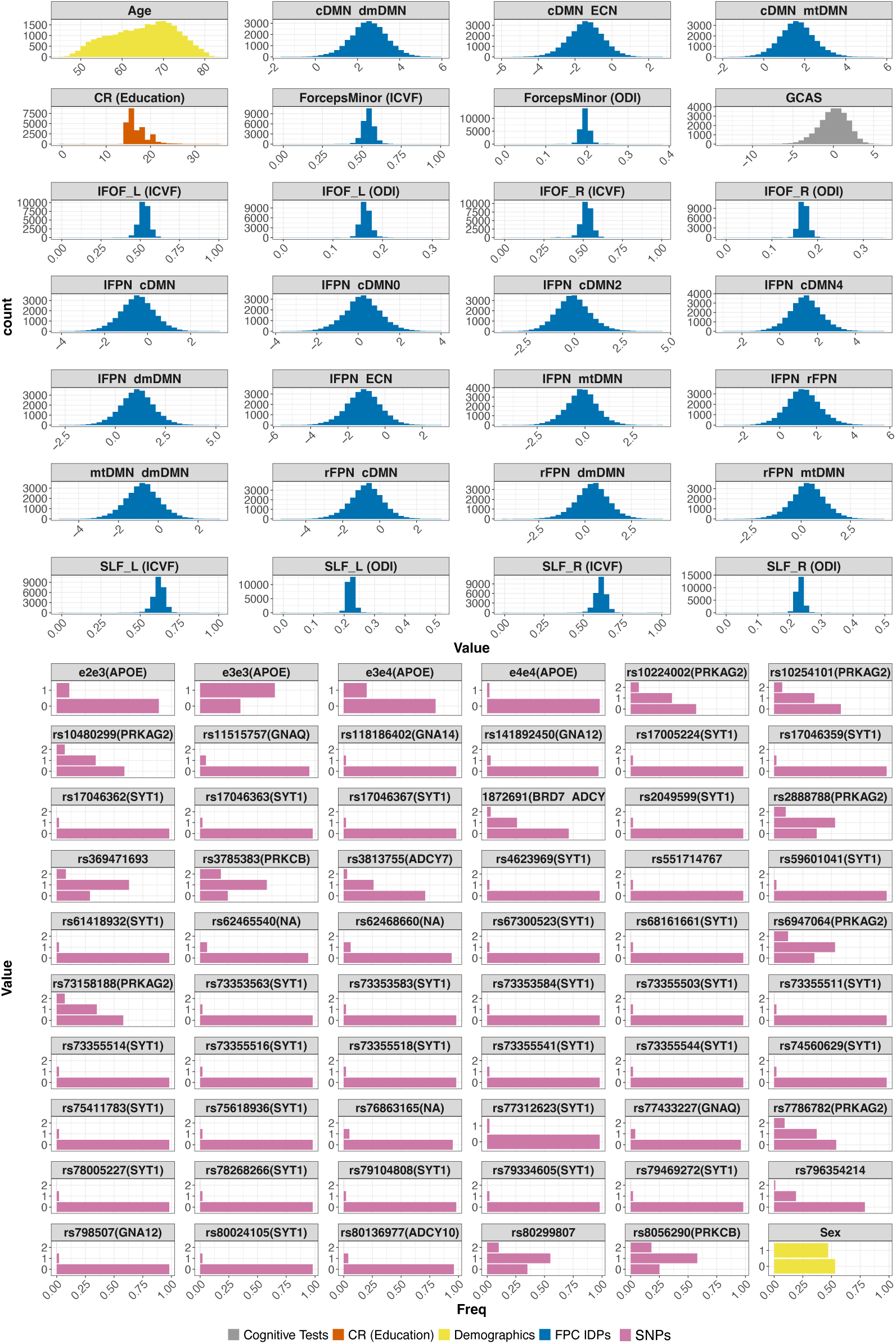
Continous and categorical distribution of selected features with information on cognitive tests, demographics, education, genes and multimodal neuroimaging data.

**Supplementary Figure 6.**
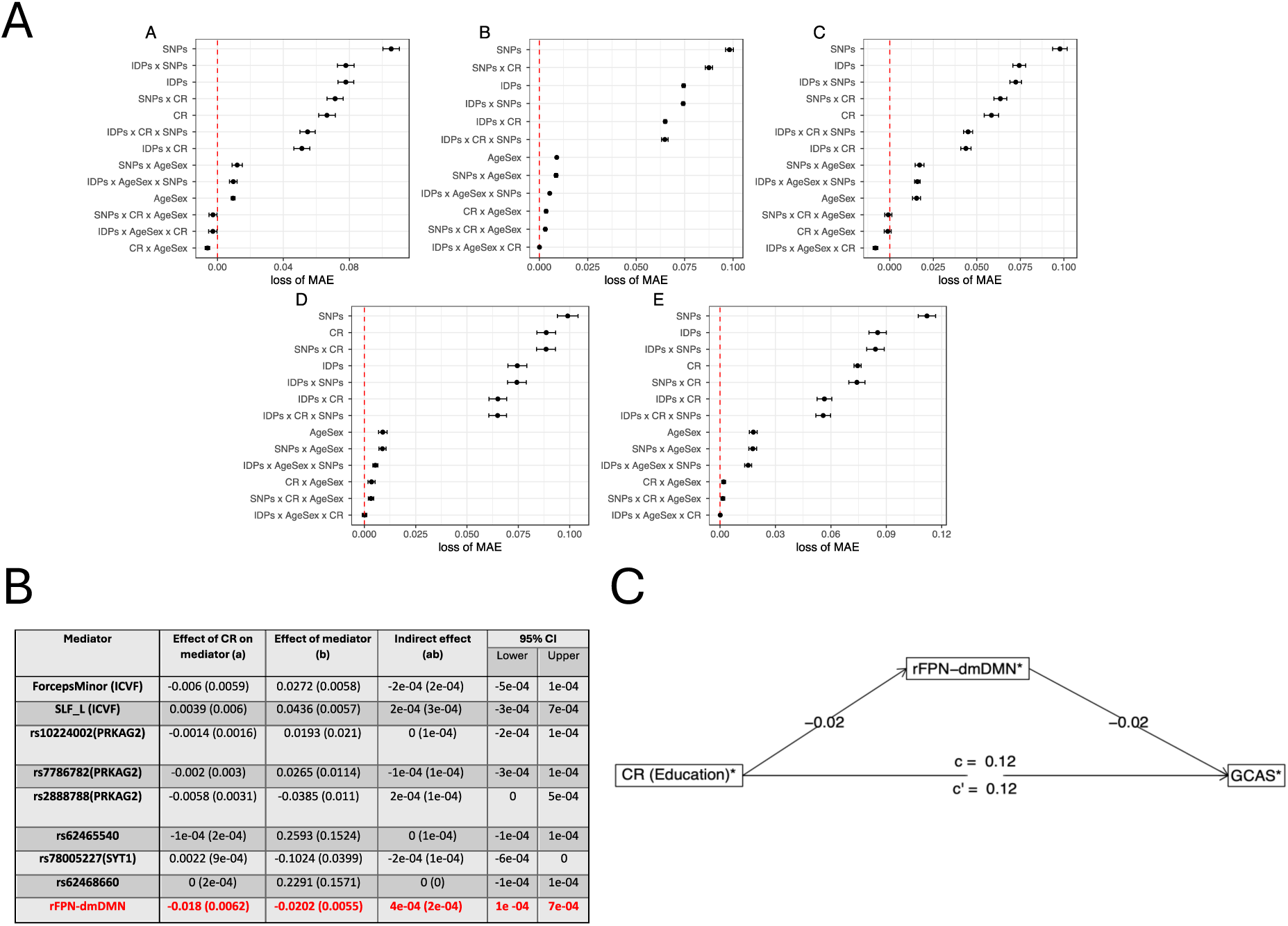
Imputed Dataset (K-nearest neighbours using 3 neighbours in tidymodels) given that only 20.2% of our dataset is missing but 54% (14,151/26,227) of participants are deleted from our analysis in complete case analysis, we performed a sensistivity analysis with imputed data to understand how and if results differed substantially with more data. A) Subsequently using imputation data, same analaysis was performed and the combination of IDPs, AgeSex and CR came as best performing too in most algorithms as is the case for the complete case analysis (as in Figure 2). Subsequently, feature importance was assessed in the same way as in complete case analysis for top performing features. Please note, 17 features differed (8 SNPs only selected in imputed data analysis and 9 IDPs and SNPs only found in complet-case analysis, with 14 chosen in common). B) Mediation results using for top 9 most frequent important variables. C) In mediation analysis, rFPN-dmDMN connectivity was found as a significant in both complete case analysis and imputed data analysis ascertaining the robustness of findings.

## Appendix

**UK Biobank ID Fields used**

**Demographic information and cognitive data**

f.34.0.0:Clinical_Birth_Year

f.52.0.0:Clinical_Birth_Month

f.53.2.0:Clinical_Date_Assessment

f.31.0.0:Clinical_Sex

f.845.0.0:Education_0

f.845.1.0:Education_1

f.845.2.0:Education_2

f.20023.2.0:Cog_ReactionTime

f.4282.2.0:Cog_NumericMemory

f.20016.2.0:Cog_FluidIntelligence

f.6348.2.0:Cog_TrailA

f.6350.2.0:Cog_TrailB

f.6373.2.0:Cog_MatrixPatterns

f.21004.2.0:Cog_TowerRearr

f.23324.2.0:Cog_SymbolSubs

f.20197.2.0:Cog_PairedAssoc

f.399.2.2:Cog_PairMatch

**MRI data**

f.25661.2.0:WM_ICVF_ForcepsMinor

f.25662.2.0:WM_ICVF_IFOF_Left

f.25663.2.0:WM_ICVF_IFOF_Right

f.25671.2.0:WM_ICVF_SLF_Left

f.25672.2.0:WM_ICVF_SLF_Right

f.25688.2.0:WM_OD_ForcepsMinor

f.25689.2.0:WM_OD_IFOF_Left

f.25690.2.0:WM_OD_IFOF_Right

f.25698.2.0:WM_OD_SLF_Left

f.25699.2.0:WM_OD_SLF_Right

The 25 components group-ICA spatial maps and connection edges are available online (http://www.fmrib.ox.ac.uk/ukbiobank).

c7, c11, c15, c16, c20, c21, c29, c33, c34, c35, c106, c110, c111, c112,c114

